# A multi-institutional study to investigate the sparing effect after whole brain electron FLASH in mice: Reproducibility and temporal evolution of functional, electrophysiological, and neurogenic endpoints

**DOI:** 10.1101/2024.01.25.577164

**Authors:** Olivia GG Drayson, Stavros Melemenidis, Nikita Katila, Vignesh Viswanathan, Enikö A Kramár, Richard Zhang, Rachel Kim, Ning Ru, Benoit Petit, Suparna Dutt, Rakesh Manjappa, M. Ramish Ashraf, Brianna Lau, Luis Soto, Lawrie Skinner, Amu S. Yu, Murat Surucu, Peter Maxim, Paola Zebadua-Ballasteros, Marcelo Wood, Janet E. Baulch, Marie-Catherine Vozenin, Billy W Loo, Charles L. Limoli

## Abstract

**Purpose:** Ultra-high dose-rate radiotherapy (FLASH) has been shown to mitigate normal tissue toxicities associated with conventional dose rate radiotherapy (CONV) without compromising tumor killing in preclinical models. A prominent challenge in preclinical radiation research, including FLASH, is validating both the physical dosimetry and the biological effects across multiple institutions.

**Methods:** We previously demonstrated dosimetric reproducibility of two different electron FLASH devices at separate institutions using standardized phantoms and dosimeters. In this study, we compared the outcome of FLASH and CONV 10 Gy whole brain irradiation on female adult mice at both institutions to evaluate the reproducibility and temporal evolution of multiple endpoints.

**Results:** FLASH sparing of behavioral performance on novel object recognition (4 months post-irradiation) and electrophysiologic long-term potentiation (LTP, 5-months post-irradiation) was reproduced between institutions. Interestingly, differences between FLASH and CONV on the endpoints of hippocampal neurogenesis (Sox2, doublecortin), neuroinflammation (microglial activation), and electrophysiology (LTP) at late times were not observed at early times.

**Conclusions:** In summary, we demonstrated reproducible FLASH sparing effects between two beams and two institutions with validated dosimetry. FLASH sparing effects on the endpoints evaluated manifested at late but not early time points.

## Introduction

In rodent models, ultra-high dose rate FLASH has been shown repeatedly to spare a number of sequelae associated with radiation injury to normal tissue while maintaining uncompromised tumoricidal efficacy compared to the same doses of conventional dose rate irradiation (CONV) (1,2). Studies of whole-brain FLASH in mice have demonstrated sparing of long-term cognitive function evaluated with a multitude of neurobehavioral assessments, as opposed to the significant impairments found routinely in animals exposed to CONV (3–9). Behavioral outcomes correspond to similar sparing effects observed on electrophysiological assessments of long-term potentiation (LTP), protection of the cerebral vasculature and neuronal morphology along with reductions in neuroinflammation (5,6,8–10).

One challenge that confronts researchers in preclinical FLASH radiobiology involves reproducing consistent beam parameters with validated dosimetry across institutional sites, and translating these beam delivery modalities to reproducible biological results (11). Characterizing carefully validated physical parameters is a prerequisite for obtaining reproducible and robust biological effects. A cross-institutional dosimetric comparison study was previously conducted between a FLASH-Trilogy and the eRT6 using a standardized phantom, auditing different electron FLASH irradiation platforms and configurations (12). More recently, another cross-institutional dosimetric comparison was conducted across three institutes using a 3-D printed anatomically realistic mouse phantom (13). In parallel, a dosimetric and biological cross-institutional comparison was performed between the eRT6 and the proton-FLASH beam at the Paul Scherrer Institute (PSI) (14). In all studies, the ability to replicate consistent and accurate delivery of FLASH and CONV doses was replicated across all institutions.

With cross-validated physical dosimetry at multiple institutions, this study aimed to confirm the biological equivalence of electron FLASH beams with distinct temporal structure. Animals were irradiated at both institutions replicating as closely as possible the same dose, dosing regimen, and treatment field in FLASH and CONV, with centralized assessment of neurological outcomes at a single independent reference center.

For endpoints, we selected two functional outcomes in the CNS that have proven reliable indicators of the FLASH effect. Neurocognitive function remains a gold standard for CNS outcomes and the FLASH effect, and when coupled with assessments of long-term potentiation, a readout of synaptic plasticity, provides two independent and unequivocal markers of neurological sparing. However, we also sought to complement these studies in efforts to assess the early radiation response of the brain to dose rate modulation. Thus, in a secondary goal, we sought to link early (<1 month) radiation-induced sequelae in the brain to the adverse late functional outcomes that take many months to manifest. Early studies finding cyclical waves of secondary reactive mediators involving reactive oxygen and nitrogen species and inflammatory molecules gave rise to the concept that the irradiated brain may never return to baseline (15,16). Based on the foregoing, we sought to evaluate whether a similar approach could help identify early mediators of the FLASH effect in the brain by analyzing selected functional, inflammatory, and neurogenic outcomes at times up to three weeks post-irradiation. Here we provide comparative biological outcomes obtained between two institutions delivering physically cross-validated CONV and FLASH electron beams and describe the disconnect that remains when trying to visualize a possible continuum between early effects and the manifestation of late effects in the irradiated brain.

## Materials and Methods

### Animals

Female C57BL/6J mice (*n*=16 animals per group) were purchased from Jackson Laboratories (Sacramento, CA) and allowed to acclimate. Mice were 11-12 weeks of age at time of irradiation. Animal procedures were conducted in accordance with NIH guidelines and Institutional Animal Care and Use Committees (IACUCs: APLAC-27939, AUP-21-025) for animal experimentation.

A separate cohort of female C57BL/6J mice (*n*=8-16 animals per group) were purchased from Charles River Laboratories (France) and allowed to acclimate. Mice were 12 weeks of age at time of irradiation. Animal procedures were conducted in accordance with the ethics committees (VD2920, VD3241, VD3603 and AUP-21-025) for animal experimentation.

All studies were conducted with tumor-free animals to avoid confounding effects of CNS tumors on cognition and associated pathology.

### Irradiation

Irradiation was performed at two institutions, with two different electron linear accelerators (linac) (12). Comparative phantom dosimetry had been conducted as previously published (13). The irradiation field was matched between institutions by using in both cases a collimator with a circular aperture of 1.7 cm placed in contact with the dorsal surface of the mouse head, with the rostral border just caudal to the eyes such that the whole encephalon region was irradiated while limiting irradiation of the eyes, mouth, and the rest of the body.

Irradiations at one institute were performed using a Varian medical linac (Trilogy, Varian Medical Systems, Inc., Palo Alto) as described previously (17). Mice received 10 Gy whole-brain irradiation delivered in a single fraction as either CONV (0.10 Gy/s mean dose rate) or FLASH delivered in 5 pulses (225 Gy/s mean dose rate). Details of the irradiation parameters can be found in **Supplementary Table 1**. Whole brain irradiation (WBI) was performed under isoflurane anesthesia (3% for induction, 2% for maintenance, in clinical room air at 2 L/min). The total duration of anesthesia was matched for all groups at 7 minutes total duration (FLASH, CONV, and unirradiated control). Each mouse was irradiated within a stereotactic supine positioner for consistent positioning relative to the circular collimator, with the entrance surface at a distance of 18.7 cm from the electron scattering foil (**Supplementary Figure 1**). The absorbed dose and spatial profiles were measured using radiochromic films positioned at the entrance surface of the mouse, the results of which are presented in **Supplementary Figure 2**.

Irradiation at the other institute were performed using a prototype 6MeV Oriatron 6e electron beam linear accelerator (PMB-Alcen, France) as described previously (7). Physical dosimetry of this linac has been extensively described in previous publications (18,19). Mice also received 10 Gy WBI delivered in a single fraction as either CONV (0.1 Gy/s) or FLASH delivered in a single (1.8 microsec) pulse (5.5 10^6^ Gy/s). Details of the irradiation parameters have been published previously (7). WBI was performed under isoflurane anesthesia in air.

### Transportation

Following irradiations at each institute, mice were returned to their standard housing environment and monitored daily for body weight, appearance, and respiratory rate for the first week and every two days thereafter. Three weeks after irradiation, in accordance with Institutional Animal Care external transportation protocols, the mice were transported to the centralized reference center where they acclimated until behavioral, electrophysiological, and/or metabolic assays were performed.

### Cognitive testing

#### Novel Object Recognition (NOR)

The NOR task was conducted four months after irradiation in a dimly lit room inside a 30×30×30 cm arena lined with a layer of fresh corncob bedding. The arena walls are cleaned after each trial with 10% alcohol and the plastic toys are cleaned after each round with 70%. An overhead camera records habituation, training, and testing for both manual and automated analyses.

Mice were handled for 2 minutes a day for 4 days and habituated to the empty arena for 5 minutes a day for 4 days, beginning on the third day of handling. 24-hours later mice were trained in the arena with two identical plastic toys for 5 minutes before being returned to their cages. After a 5-minute consolidation period, mice were returned to the arenas with one of the original toys having been replaced with an object differing in color and shape. They were able to explore the test configuration for 5 minutes and then returned to their cages. This is the short-term version of the NOR test as the mice have only a 5-minute gap between training and testing. Cognitive testing was performed following previously published protocols (8,20,21).

Cognitive function is determined by calculating the Discrimination Index (DI) as shown in **Equation 1**. Animals have a tendency to explore novel objects and therefore the cognitively impaired animals should show a lower DI than a healthy control. The NOR task is reliant on intact perihinal cortex function (15(22,23)

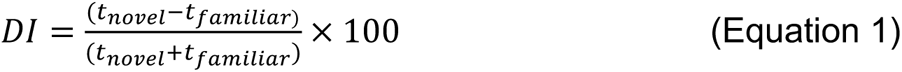

### Electrophysiology

After completion of behavioral testing, a subset of the cohort (n=6/treatment/institution) was sacrificed for electrophysiology. The uteri of female mice were dissected and weighed prior to LTP assessments, confirming that none of the subjects were in estrus. The preparation of the hippocampal slices has been described previously (24). Anesthetized mice were decapitated, and the brains were quickly transferred to an ice-cold oxygenated dissection medium containing (in mM): 124 NaCl, 3 KCl, 1.25 KH_2_PO_4_, 5 MgSO_4_, 0 CaCl_2_, 26 NaHCO_3_, and 10 glucose. Hippocampal slices (340 μm, coronal) were cut from a vibratome (Leica, Model:VT1000S) before transfer to an interface recording containing prewarmed (31 ± 10°C) artificial cerebrospinal fluid (aCSF) composed of (in mM): 124 NaCl, 3 KCl, 1.25 KH_2_PO_4_, 1.5 MgSO_4_, 2.5 CaCl_2_, 26 NaHCO_3_, and 10 glucose. The slices were exposed to warm, humidified 95% O_2_/5% CO_2_ while being continuously perfused at a rate of 1.75-2 mL/min. Recordings were initiated after a minimum of 2 hours of incubation.

Field excitatory postsynaptic potentials (fEPSPs) were recorded from CA1b stratum radiatum apical dendrites using a glass pipette filled with 2M NaCl (2-3 MΩ) in response to orthodromic stimulation (twisted nichrome wire, 65 μm diameter) of Schaffer collateral-commissural projections in CA1 stratum radiatum. Pulses were administered 0.05 Hz using a current that elicited a 50% maximal spike-free response. After maintaining a stable baseline (20 minutes), LTP was induced by delivering 5 ‘theta’ bursts, with each burst consisting of four pulses at 100 Hz separated by 200 milliseconds (*i.e.*, theta burst stimulation or TBS). The stimulation intensity was not increased during TBS. Data were collected and digitized by NAC 2.0 (Neurodata Acquisition System, Theta Burst Corp., Irvine, CA) and stored on a disk. The fEPSP slope was measured at 10–90% fall of the slope and data in figures on LTP were normalized to the last 20 minutes of baseline.

### Immunohistochemistry

Immunohistochemistry was performed on brains harvested two days after irradiation (from a single institute) for microglia (CD68, IBA-1) and neural stem cells (SOX2), and 1 week, 2 weeks, and 3 weeks for immature neurons (doublecortin, DCX). Animals were deeply anesthetized with isoflurane and euthanized with saline with heparin (10 U/mL, Sigma-Aldrich) and then 4% paraformaldehyde through intracardiac perfusion. The brains were cryoprotected using a sucrose gradient (10%, 20%, and 30%) and then coronally sectioned (30 μm thick) using a cryostat (Leica Microsystems, Germany). Three sections around the midpoint of the hippocampus were selected per animal for each stain. For DCX immunostaining, every seventh section of the right hemisphere of the brain through the rostrocaudal span of the mid-hippocampus (4 sections) were stained and included in the analysis. The sections were stained free-floating and then mounted onto slides for imaging.

For the immunofluorescence analysis of microglia, the following primary and secondary antibodies were used: rabbi anti-IBA-1 (1:500, Wako), rat anti-mouse CD68 (1:500 BioRad), goat anti-rabbit 488 (1:500, Invitrogen), goat anti-rat 647 (1:1000, Abcam), and DAPI nuclear counterstain (Sigma-Aldrich). IBA-1 positive cells were visualized under fluorescence as green, CD68 as red and nuclei as blue. For the immunofluorescence analysis of neural stem cells, SOX2, an HMG box transcription factor, is stained using the following primary and secondary antibodies: goat anti-SOX2 polyclonal (1:100, Novus), donkey anti-goat 488 (1:200, Invitrogen), and the same DAPI counterstain (half the concentration used in IBA-1/CD68 stain). SOX2 positive cells fluoresce green, and nuclei blue. For the Doublecortin (DCX) immunofluorescence stain, the following antibodies were used: rabbit anti-DCX (1:750, abcam), donkey anti-rabbit AF488 (1:500, abcam), and the DAPI nuclear stain. DCX positive cells fluoresce green and DAPI stained nuclei fluoresce blue.

Immunofluorescent sections were imaged using a Nikon Eclipse Ti C2 microscope to obtain 20 to 30 z-stacks (1024×1024 pixels, 1μm each) using 40× magnification PlanApo oil-immersion lens (Nikon). For quantification of IBA-1 and CD68 positive cells, 3D deconvolution and surface reconstruction were carried out using Imaris software (v8.0, Bit Plane Inc., Switzerland), and the overlapped volume between IBA-1 positive cells and CD68 positive cells was quantified. For quantification of SOX2, the Imaris cell tool was used to count positive cells located only in the Subgranular Zone (SGZ). For DCX immunofluorescently stained sections, images were obtained using a Leica Dmi8 microscope. Images from 12-15 z-stacks (1024×1024 pixels, 0.5μm each) using 10× magnification were obtained and post-processed with extended depth of field. The images were then quantified using ImageJ software to manually count the number of DCX-positive neurons in the granular cell layer-subgranular zone (GCL-SGZ) and represented as the sum of the DCX-positive neurons in the GCL-SGZ.

### Statistical Analysis

All statistical analyses were performed using Prism software (Graphpad Software Inc, San Diego California, USA, version 9.4.1). One-way analysis of variance (ANOVA) was used to assess significance between treatment groups. Where overall group effects were significant, Bonferroni’s post-hoc multiple comparison test was performed to compare the unirradiated control and FLASH groups against the CONV cohort. Data are presented as Mean ± SEM and any values of *P*≤0.05 were considered statistically significant.

## Results

### FLASH sparing is replicated between institutions at late timepoints

#### Cognitive function is spared in FLASH cohorts from both institutions

We investigated the impact of electron FLASH and CONV whole brain irradiation delivered with FLASH-trilogy on cognitive function in comparison to previous results obtained after electron FLASH and CONV WBI with eRT6. CONV irradiation is known to induce impairments in learning and memory, effects that FLASH has been shown to minimize. (6,8,9,20).

Four months after irradiation, animals from institute 1 underwent behavioral testing. Animals receiving 10 Gy FLASH WBI were statistically indistinguishable from unirradiated controls in the Novel Object Recognition (NOR) task, whereas animals receiving 10 Gy CONV WBI had a statistically significant reduction in their Discrimination Index (DI) (**Fig 1A**, one-way ANOVA: F(2,43)=5.1, *P*=0.0096).

**Figure 1:**
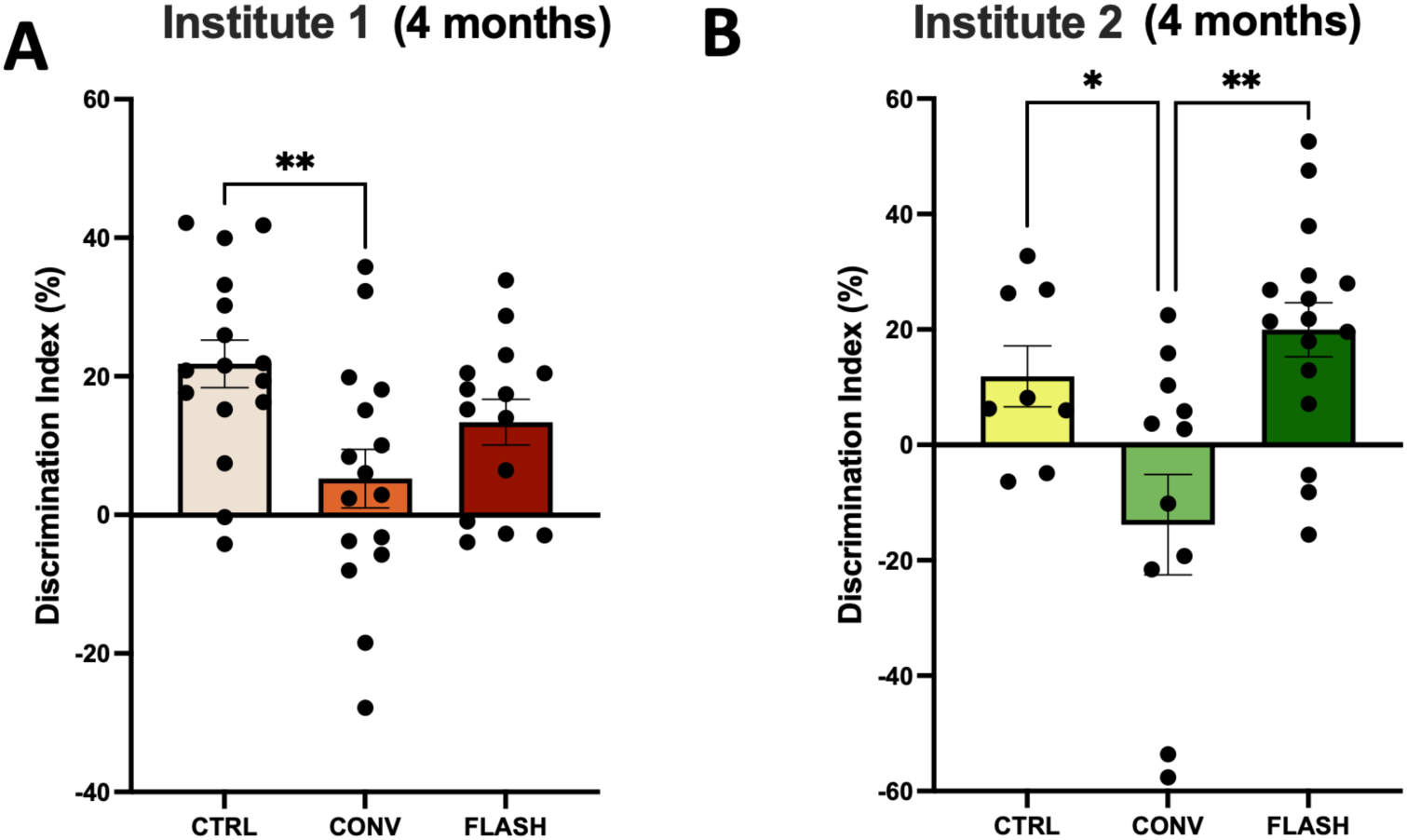
FLASH sparing of performance on the Novel Object Recognition (NOR) test late (4 months) after 10 Gy single fraction whole brain irradiation is reproduced between institutions. A) NOR scores from Institute 1 were scored manually by core facility experts. CONV irradiated animals had significantly lower discrimination index than unirradiated controls, but FLASH irradiated animals were not statistically different from unirradiated controls. B) Previously published (6) NOR scores from Institute 2 exhibit similar effects between cohorts where significance a significance decrease in DI was seen in the CONV irradiated animals compared to unirradiated controls, and the FLASH irradiated animals had significantly higher DI values compared to the CONV irradiated animals. Data were analyzed using one-way ANOVA and the Bonferroni multiple comparisons test (n=16/group). *,P σ; 0.05; **, P σ; 0.01; ns, not significant.

This result matches previous data from the NOR behavioral task conducted on mice 4 months after irradiation at Institute 2 with one fraction of 10 Gy WBI (6). A statistically significant difference was observed between the CONV cohort and unirradiated controls and between CONV and FLASH cohorts **(Fig 1B**, one-way ANOVA: F(2,33)=7.944, *P*=0.0015).

#### Electrophysiological evaluation shows preservation of Long-Term Potentiation (LTP) by FLASH in both cohorts

Two weeks after completion of behavioral testing, Theta Burst Stimulation (TBS) was applied to hippocampal slices to induce LTP. Five theta bursts to Schaffer collaterals induced fEPSP (**Fig. 2**). After this initial short-term potentiation, a gradual decay in the fEPSP was observed in all cohorts at both institutes as shown in **Figures 2 A and B**. The stable potentiation value in the fEPSP slope was significantly lower in CONV groups than unirradiated controls but not in FLASH groups. This result is quantified by mean potentiation measured 50-60 minutes after TBS (**Fig. 2**). The mean potentiation results showed the same effect in CONV cohorts with the same level of significance. Institute 1 cohort one-way ANOVA: F(2,30)=54.39, *P*<0.0001, Bonferroni *post-hoc*: unirradiated control vs CONV: *P*<0.0001; FLASH vs CONV: *P*<0.0001. Institute 2 cohort one-way ANOVA: F(2,33)=30.11, *P*<0.0001, Bonferroni *post-hoc*: unirradiated control vs CONV: *P*<0.0001; FLASH vs CONV: *P*<0.0001.

**Figure 2:**
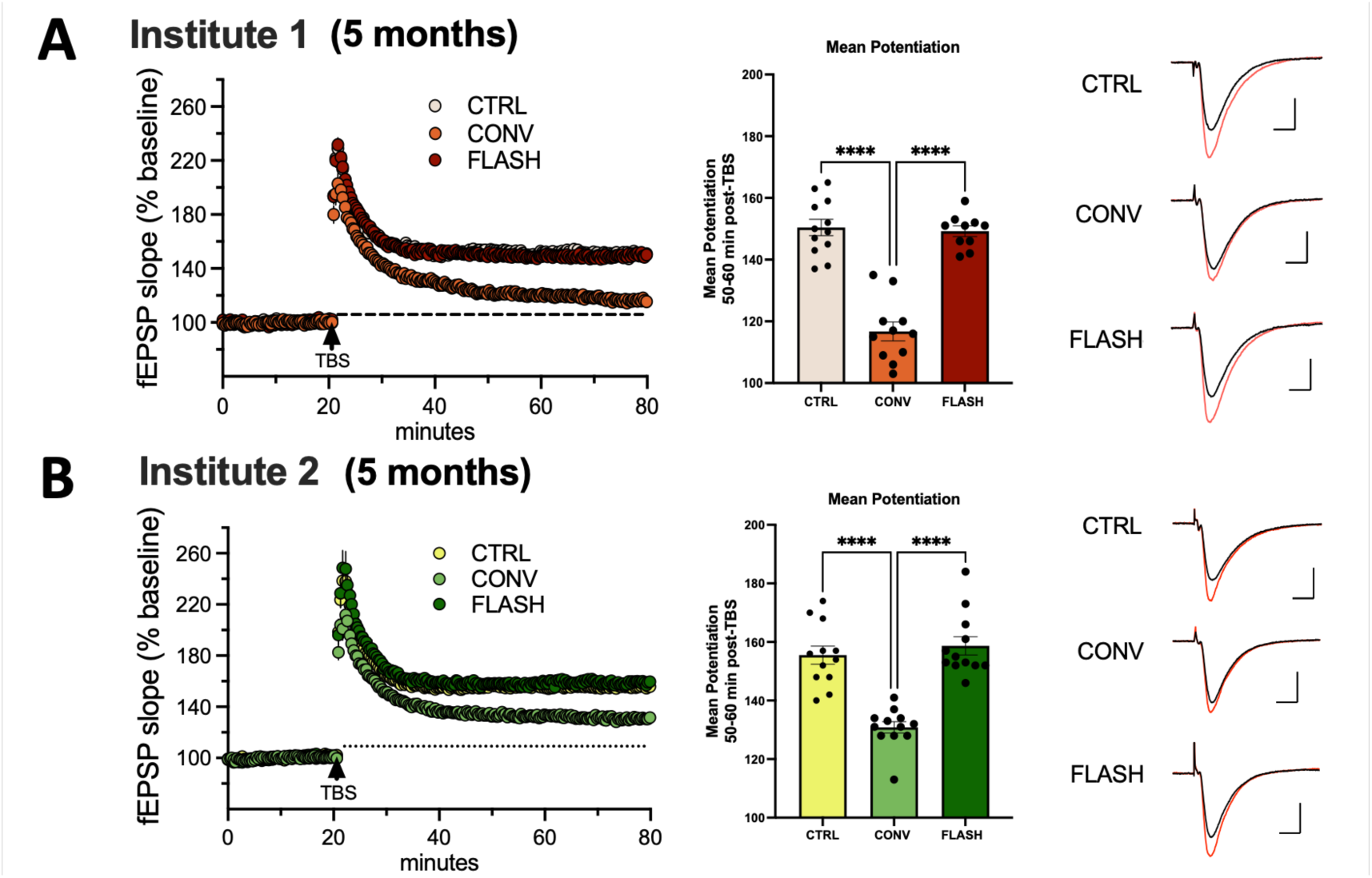
FLASH sparing of Long Term Potentiation (LTP) in late (5 month) mice after 10 Gy single fraction whole brain irradiation is reproduced between institutions. A) The LTP measurement results from Institute 1. (left) Following a stable 20 min baseline recording, the slope of the field Excitatory Postsynaptic Potential (fEPSP) as a percentage of baseline shows an immediate increase in potentiation after delivering Theta Burst Stimulation (TBS). The combined slope of the CONV irradiated cohort fails to stabilize unlike the unirradiated control or the FLASH irradiated group. (middle) The mean potentiation 50-60 minutes post-TBS for each treatment group. The mean potentiation is significantly lower in the CONV group compared to both the unirradiated controls and the FLASH group. (right) Representative traces collected during baseline (black line) and 50-60 min post-TBS (red line) for each group. Scale = 1mV/5ms. B) The fEPSP slope as percentage of baseline, mean potentiation and electrophysiological traces for the cohorts irradiated at Institute 2. The group differences and levels of significance are the same between each institute. Data were analyzed using one-way ANOVA and the Bonferroni multiple comparisons test (n=10-11 slices/group). ****, P ≤ 0.0001.

These functional readouts found in both irradiated cohorts at each institute demonstrate the robustness of the electrophysiological evaluation of LTP in measuring the sparing effect of FLASH and the equivalence of the FLASH irradiation generated with different irradiation devices validated at the dosimetry level and located in two different institutions.

### Timing is everything: Differential effects of FLASH on late CNS endpoints are not observed at acute time points

#### Electrophysiological evaluation shows no impact of irradiation on LTP 2 weeks post-irradiation

For selected CNS endpoints, much less is known of how FLASH might differentially impact acute responses. Measurements of LTP as described above were repeated on 10-week-old mice 2 weeks after 10 Gy CONV and FLASH irradiation. Interestingly, and in marked contrast to our past publications conducted at > 4 month post-irradiation, there was no observable difference between the irradiation groups in mean potentiation at an early time point (**Fig. 3**, one-way ANOVA: F(2,31)=0.3822, *P*=0.6856). There also was no deviation in the fEPSP slope from the unirradiated control group for either FLASH or CONV treatment groups as shown in **Figure 3**.

**Figure 3:**
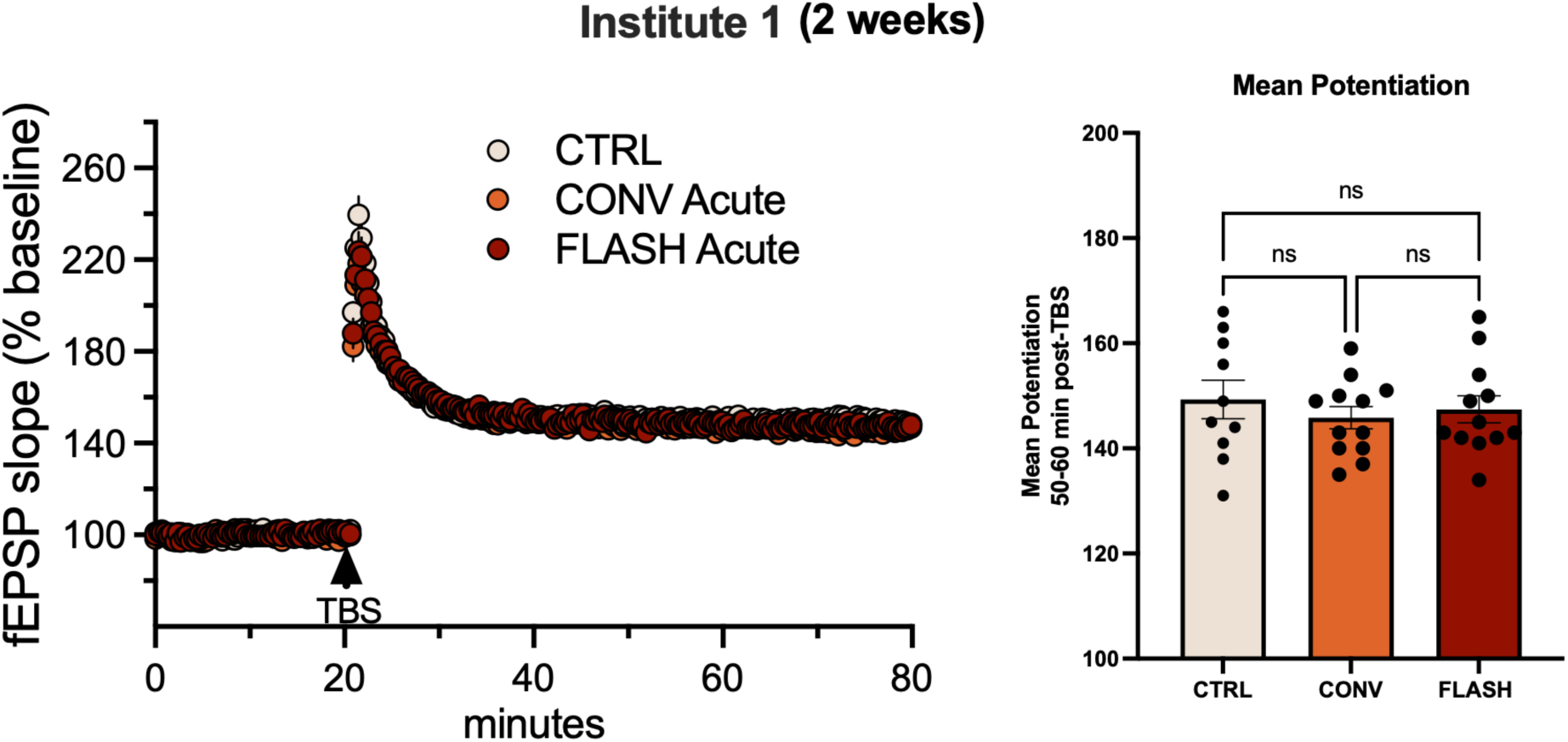
LTP is not altered acutely (2 weeks) after 10 Gy single fraction whole brain irradiation by either FLASH or CONV. The fEPSP slope as percentage of baseline is indistinguishable between unirradiated controls, CONV, and FLASH groups irradiated at Institute 1. No significant difference between mean potentiation 50-60 min post-TBS was observed in either irradiation group. Data were analyzed using one-way ANOVA and the Bonferroni multiple comparisons test (n=10-12 slices/group). ns, not significant. This contrasts with the decrement in LTP observed to emerge later after CONV but not FLASH (see Fig 2).

#### Neuroinflammation at an acute timepoint is equally elevated in FLASH and CONV irradiated mouse brain

Immunohistochemical staining for the IBA-1 microglial marker and CD-68 marker for reactive microglia indicated a significant increase in neuroinflammation 48 hours after irradiation. However, at this acute timepoint both FLASH and CONV increased levels of reactive microglia to similar extents compared to the unirradiated control group (**Fig. 4**, one-way ANOVA: F(2,71)=4.964, P=0.0096). Previous studies have shown CONV irradiation to trigger a persistent elevation in neuroinflammation at both acute and chronic timepoints (25), but the marked attenuation of these effects reported long after FLASH (5,6) was not evident at acute timepoints.

**Figure 4:**
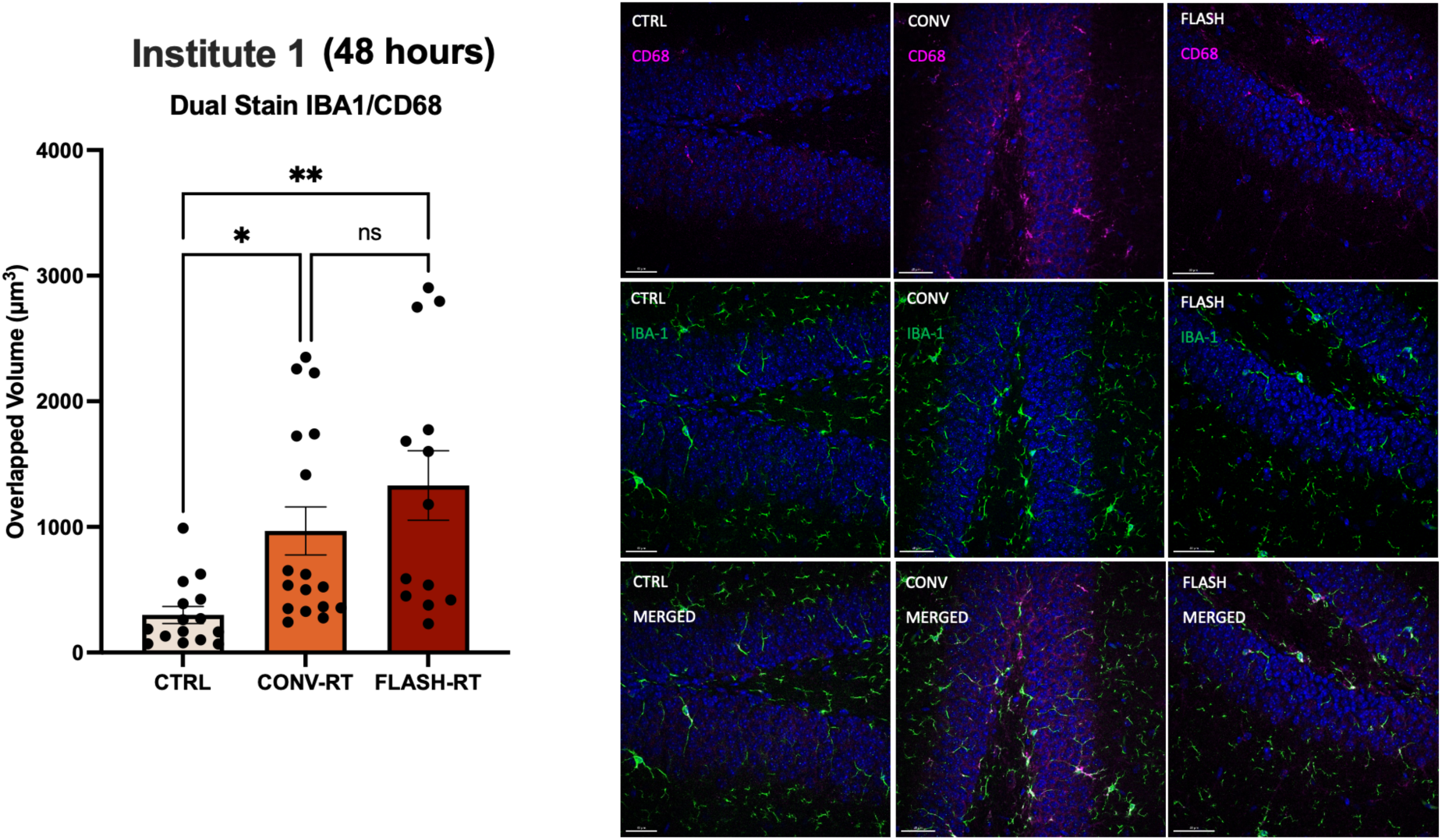
Neuroinflammation in the hippocampus increases acutely (48 hours) after 10 Gy single fraction whole brain irradiation by either FLASH or CONV. Dual Stain of IBA-1 (microglial stain) and CD68 (activated microglial stain) in the hippocampus of mice 48 hours after irradiation at Institute 1. (Left) The volume of overlap of the two stains is plotted with each data point representing a single section. A significant increase in the volume of activated microglia was observed in both the CONV and FLASH cohorts compared to unirradiated controls. (Right) Representative images of the CD68 stain alone (top) with DAPI in blue, the IBA-1 stain alone (middle) and the combined stain (bottom) for each of the three treatment groups. Data were analyzed using one-way ANOVA and the Bonferroni multiple comparisons test (n=5-11/group, each datapoint represents an average of 2-3 sections/animal). **, P ≤ 0.01; ns, not significant. This is in contrast to the resolution of neuroinflammation observed to emerge later after FLASH but not CONV.

#### Initial response of neurogenic populations is similar between FLASH and CONV but diverges with time

In the same cohort of animals from institute 1, Sox2 immunohistochemical staining was conducted to evaluate the impact of 10 Gy of cranial irradiation on the population of neural progenitor cells at the acute 48 hour post-irradiation time point. The transcription factor Sox2 is expressed in radial glial-like neural stem/progenitor cells and is essential for self-renewal and differentiation (26). Only Sox2+ cells in the SGZ were counted. The Sox2 stain revealed no significant difference between either the CONV or FLASH irradiated cohorts relative to the unirradiated controls 48 hours post-irradiation (**Fig. 5**) (F(2,9)=0.5034, *P*=0.6205), suggesting no significant loss of neural stem cells or neural progenitor cells in the dentate gyrus at 48 hours post-irradiation.

**Figure 5:**
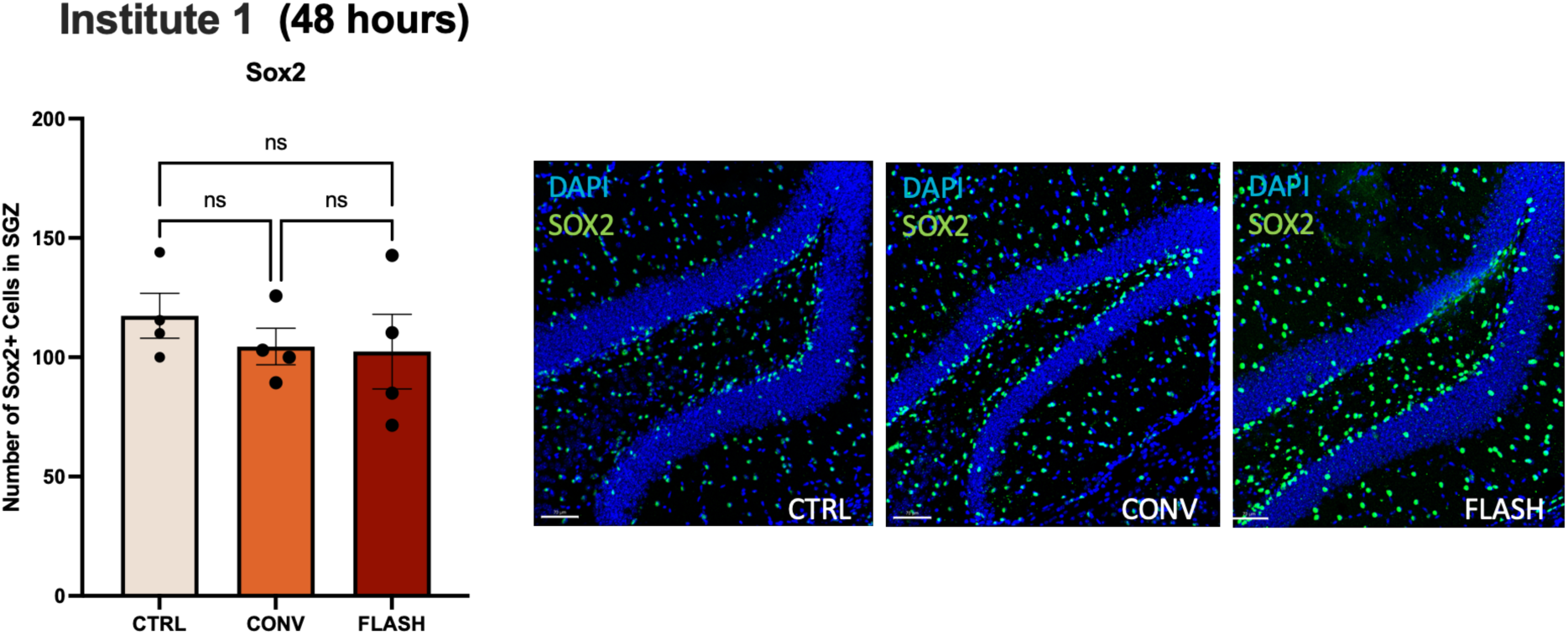
The number of neural stem cells in the hippocampus is unchanged acutely (48 hours) after 10 Gy single fraction whole brain irradiation by either FLASH or CONV. Sox2 stain of the SGZ of the hippocampus of mice 48 hours after irradiation of 10Gy at Institute 1. No significant difference between any irradiation group was observed.

The DCX stain was performed at one week, two weeks and three weeks post-irradiation to evaluate the acute impact on immature neurons. DCX is often used as a surrogate marker for neurogenesis (27). The number of DCX+ cells was severely reduced in both irradiated groups compared to unirradiated controls at one and two weeks (**Fig. 6**) F(2,8)=195.8, *P*<0.0001 for the one week timepoint) with no difference between FLASH and CONV. Whereas the recovery of DCX+ cells plateaued at 2 weeks after CONV, levels increased significantly after FLASH at 3 weeks post-RT (unpaired t-test between FLASH and CONV at 3 weeks, t=5.862, df=6, *P*=0.0011).

**Figure 6:**
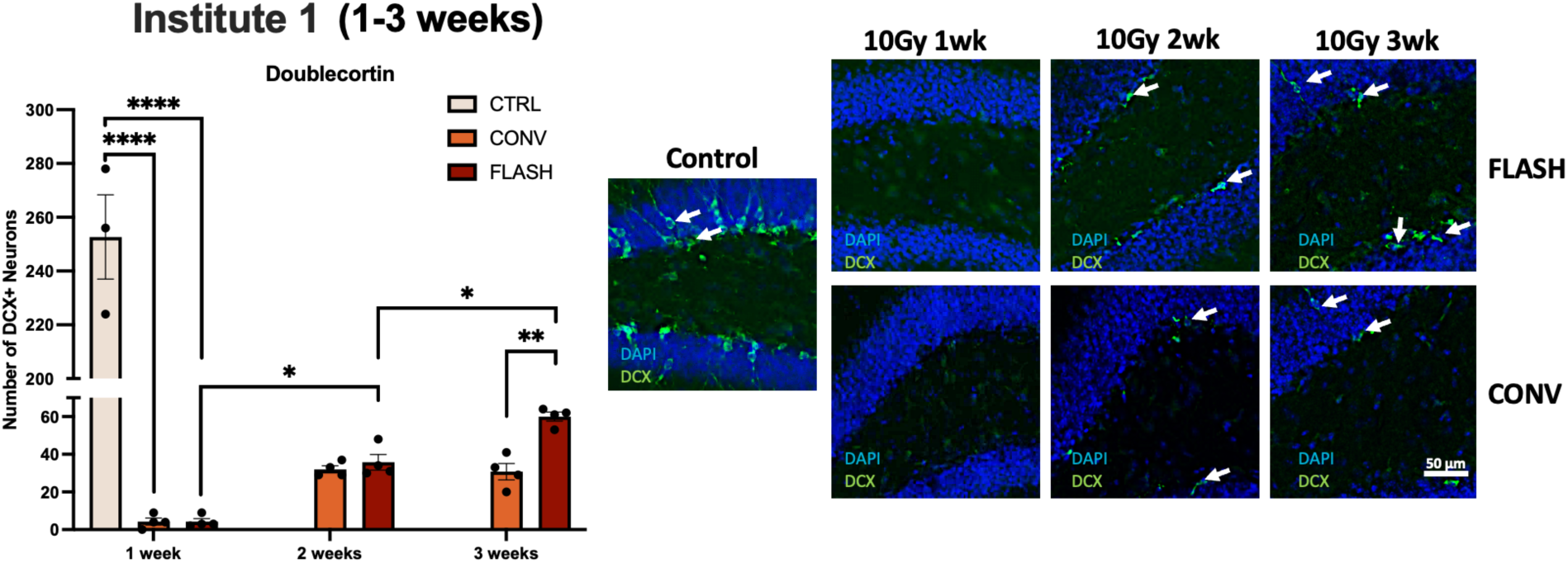
The number of immature neurons in the hippocampus is severely depleted acutely (1-2 weeks) after 10 Gy single fraction whole brain irradiation by either FLASH or CONV, but at 3 weeks recovers more with FLASH than CONV. DCX staining in the hippocampus of mice 1 week, 2 weeks and 3 weeks after irradiation of 10Gy at Institute 1. A strong depletion in DCX+ cells was observed in both CONV and FLASH groups to the same significance level at 1 week. At 2 weeks, some DCX+ cells regenerated both in FLASH and CONV. However, the recovery of the FLASH groups occurs exponentially reaching to a significant difference compared to the CONV group at 3 weeks. Data were analyzed using one-way ANOVA and the Bonferroni multiple comparisons test (n=4-5/group). *,P ≤ 0.05; **,P ≤ 0.01; ****,P ≤ 0.0001; ns, not significant.

## Discussion

The goal of this study was to validate from a biological perspective that FLASH neurological sparing after whole brain irradiation could be reproduced with different electron beams at different institutions when delivering dose and dose rates previously demonstrated to produce the FLASH effect in independent experiments. We directly compared FLASH irradiation platforms between two electron linacs that have been used extensively to examine the FLASH effect, but in this case matching the mouse model, irradiation field, and dosing regimen between them. This was followed by uniform assessment of cognition and electrophysiology at the central independent reference facility. This evaluation was preceded by a thorough dosimetric comparison of both irradiation platforms for FLASH-relevant ultra-high dose-rate and CONV dose rates (12), confirming agreement between measured and planned doses sufficient for preclinical studies such as that described in this paper. The current study confirms that equivalent biological FLASH effects are achieved by both institutions under harmonized conditions, with equivalent preservation of cognition as well as synaptic plasticity. Similar comparative studies have been conducted to evaluate the equivalence of electron and proton FLASH beams (14) and for sparing of gastrointestinal toxicities after electron FLASH (28). In the former case, neurocognitive function was spared after both FLASH modalities while tumor control and anti-tumor immunity was maintained equally between dose rates and modality. For the latter case, two electron beams were validated dosimetrically and shown to spare survival and intestinal crypt cell regeneration.

The FLASH sparing of learning, memory, attention, mood, social interaction and fear memory have been found repeatedly after electron FLASH at different doses and fractionation regimen (3,5–9). To focus our objectives, the novel object recognition task was chosen as a logical and robust endpoint to compare outcomes after CONV and FLASH between the Trilogy and eRT6 linacs. Exposure of animals to CONV from either institute resulted in cognitive impairments, with statistically significant reductions in the discrimination index observed compared to unirradiated control mice. No difference between the FLASH irradiated animals and unirradiated controls was observed, indicating that each electron linac was able to spare radiation-induced learning and memory impairment when delivering FLASH dose rates.

The prolonged sparing of learning and memory deficits after FLASH suggests a preservation of synaptic elements involved in neurotransmission. In three recent studies, hypofractionated dosing regimens (2×10 Gy, 3×10 Gy) and standard of care fractionation (10×3 Gy) was shown to preserve LTP in FLASH cohorts when assessed months after irradiation, as opposed to the significant inhibition of LTP in the CONV cohorts (8–10). Therefore, we sought to test both the irradiated cohorts institutes in the same way to identify if LTP was adversely impacted by CONV at this single dose of 10 Gy and whether FLASH could spare this detrimental effect. We found that the fESPS slope was reduced significantly for the hour post theta burst stimulation only in the CONV irradiated animals. The mean potentiation over this period was significantly inhibited in CONV cohorts from both institutions but not statistically different from unirradiated controls in FLASH cohorts from either electron linac. LTP remains a reliable standard to assess synaptic plasticity and this data further demonstrates that FLASH does not perturb the firing of Schaffer collaterals in the hippocampus, thereby preserving neurotransmission and synaptic integrity. The similarity of the LTP results between the cohorts irradiated at each institute validates the equivalence of the FLASH parameters generated by each linac and supports evidence for electrophysiological assessments as a reliable biomarker of the FLASH effect when assessed at late timepoints.

The functional equivalence of NOR and LTP outcomes between each electron linac combined with the recent article between the eRT6 and the proton beam Gantry1/PSI (14) establish certain consistent benefits of FLASH to critical functional outcomes in the CNS with various beams. However, we also sought to determine whether studies conducted at earlier times might provide more information and perhaps predictive value. Past efforts linking the onset, progression, and severity of late radiation effects in the brain to early changes in blood brain barrier permeability, apoptosis, inflammation and neurogenic cell kill have proven difficult (29,30). There has been considerable difficulty uncovering specific biomarkers of neurocognitive decline, highlighted by the rich literature derived from space radiation studies on the brain (31,32). Therefore, a secondary objective of this study was to investigate the impact of FLASH at earlier timepoints up to three weeks post-irradiation. Numerous past FLASH studies have demonstrated its ability to spare the CNS from late radiation toxicity (3,6–9,20), and the results of this study support these findings. What is much less understood is if or how FLASH might prevent early radiation responses leading to toxicity in the brain.

To investigate the potential for FLASH to modulate early toxicities in the brain, we employed one of our most reliable late markers of the FLASH effect, namely LTP. When assessed at late times (> one month) CONV irradiated cohorts exhibit significant and persistent reductions in slope of the fEPSP, effects not evident after FLASH (8–10). While this finding was replicated after a single dose of 10 Gy at five months post-irradiation, no such change was found at two weeks post-irradiation. All cohorts exhibited identical LTP firing activity, clearly indicating that the radiation-induced inhibition of LTP manifests at times later than two weeks post exposure in CONV irradiated animals, an effect that never seems to manifest in FLASH irradiated cohorts.

Elevated neuroinflammation is involved in perpetuating a host of radiation-induced and other neurological complications (37–40) and the ability of FLASH to suppress the levels of reactive microglia has proven to be another robust marker of the FLASH effect (6,8,9,33). Thus, follow-up immunohistochemical investigations were undertaken to analyze reactive microglial levels two days post-irradiation. While the levels of reactive microglia showed an increase in both irradiated cohorts, no difference between FLASH and CONV was observed. Clearly, inflammation is an immediate response of the brain to radiation damage, but the signature of radiation injury does not persist in FLASH irradiated cohorts suggesting that these incipient processes can be resolved.

Lastly, past studies have shown that FLASH can spare the neurogenic niche at late times post-irradiation (4,20), and other work has found intestinal crypt sparing after electron and proton FLASH at earlier times (28,34). Interestingly, neither FLASH nor CONV significantly impacted the number of Sox2+ cells at 48 hours post-irradiation. Data suggest that populations of early stage, transiently amplifying progenitor cells exhibit more resistance to irradiation, possibly because there was simply insufficient time to express radiation-induced lethality. In contrast, a severe depletion of DCX+ cells was induced by both CONV and FLASH one week post-irradiation compared to unirradiated controls. However, recovery of DCX+ cells began to manifest by two weeks, and by three weeks there was greater recovery of DCX+ cells in FLASH compared to CONV irradiated mice. The enhanced temporal recovery of immature neurons in the FLASH cohort is interesting and may suggest that FLASH promotes a wound healing process, although the molecular targets involved remain uncertain.

Collectively, this multicenter biological intercomparison FLASH study demonstrates that FLASH sparing of cognitive function and brain electrophysiology compared to conventional dose rate brain irradiation is a robust biological phenomenon. The use of validated dosimetry platforms across institutions provided us with a framework to perform comparable biological investigations using different beam lines. The FLASH sparing effect varied across time similarly for both institutes and was most robust for late effects such as cognition and LTP. Assessment of LTP, inflammatory and neurogenic endpoints at post-irradiation preceding three weeks yielded few dose-rate dependent changes. Data emphasize the difficulties of identifying early biomarkers of the FLASH effect in the brain but suggest that conducting longitudinal studies on more biological endpoints will be important to elucidate candidate mechanisms.

## AUTHOR CONTRIBUTIONS

All authors met the International Committee of Medical Journal Editors (ICMJE) criteria for authorship (*https://www.icmje.org/recommendations/browse/roles-and-responsibilities/defining-the-role-of-authors-and-contributors.html*). Olivia GG Drayson and Stavros Melemenidis are co-first authors, each of whom are appropriately listed first when referencing this publication on their respective CVs. Charles L. Limoli, Marie-Catherine Vozenin, and Billy W Loo Jr are co-senior/co-corresponding authors.

## DATA AVAILABILITY STATEMENT

The data that support the findings of this study are available from the corresponding author upon reasonable request.

## Supplementary Table and Figures

**Supplementary Table 1:** Number of animals and experimental beam parameters for the irradiation of mouse whole brain at Stanford University.

| **Mode** | **FLASH** | **CONV** |
| --- | --- | --- |
| **Animals per group** | 16 | 16 |
| **Beam energy [MeV]** | 16.60 | 15.73 |
| **Target dose [Gy]** | 10 | 10 |
| **Delivered dose [Gy]** | 10.08 ± 0.16 | 10.06 ± 0.36 |
| **Delivered pulses** | 5 | 9290 |
| **Pulse rate [pulses/s]** | 90 | 90 |
| **Dose per pulse [Gy]** | 2 | 1.05E-3 |
| **Dose rate [Gy/s]** | 225 | 0.10 |
| **Pulse length [μs]** | 3.75 | 3.75 |
| **Intra-pulse dose rate [Gy/μs]** | 0.53 | 2.93E-4 |

**Supplementary Figure 1:**
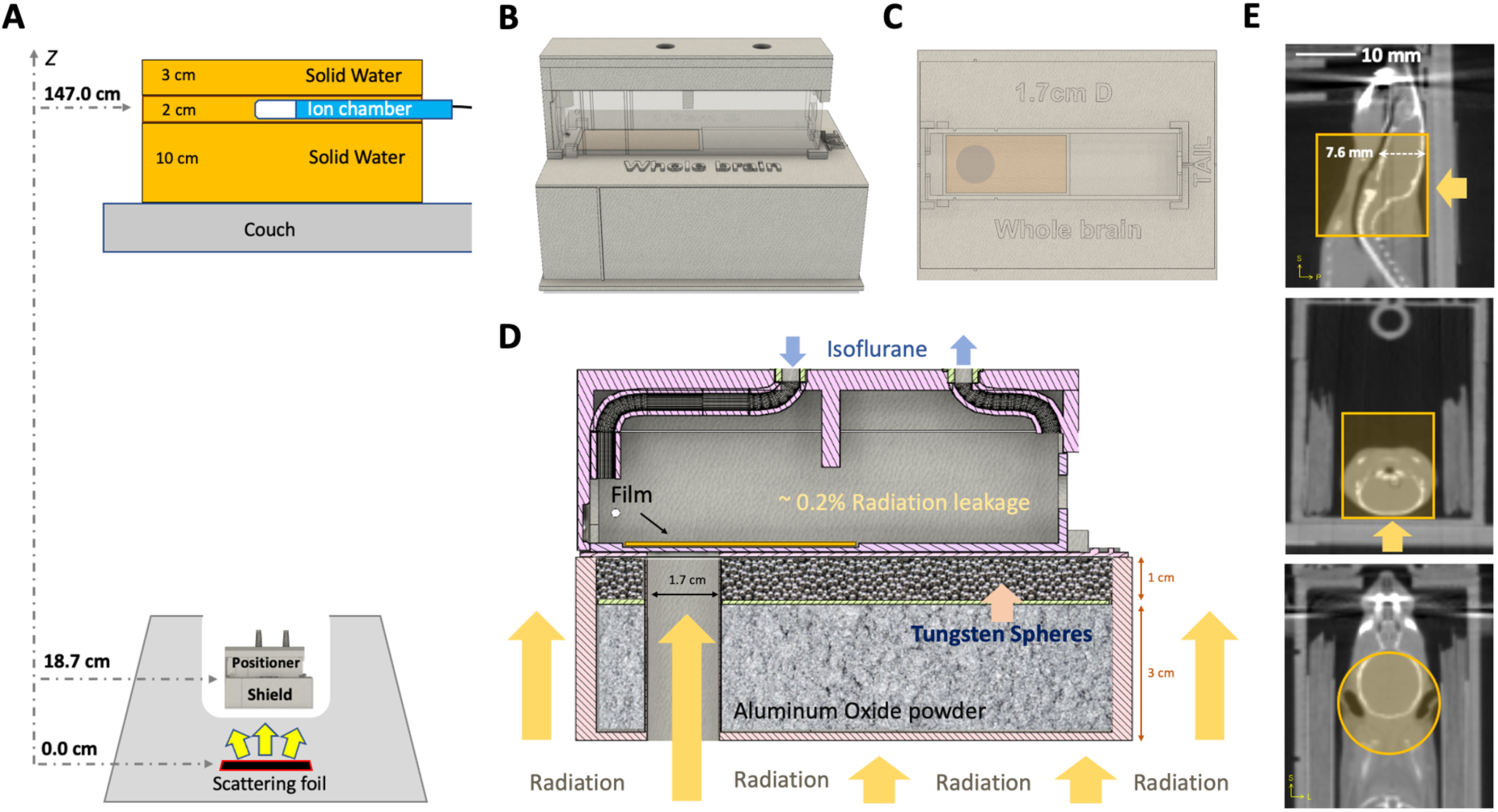
Stanford’s experimental setup with in vivo stereotactic mouse supine positioner and shield geometry. (A) Schematic diagram of the setup at Stanford for irradiations, illustrating the distance of the stereotactic mouse positioner from the scattering foil. On the top, a solid water stack encloses an ion chamber that measures the exit charge for each irradiation. (B) CAD files shows a side view of the radiation shield (bottom) with the stereotactic mouse positioner (middle; in transparency) and the isoflurane anesthesia cover (top), showing the placement of the radiochromic film used for entrance dose measurements. The vertical line on the shield indicates the middle of the radiation field and the two vertical lines on the stereotactic mouse positioner indicate the edges of the radiation field. (C) CAD file shows a top view of the shield without the anesthesia cover, illustrating the circular (1.7 cm diameter) radiation field and positioning of the radiochromic film. (D) A schematic illustration of the radiation assembly as shown in (B), using a sagittal cross section plane of the CAD file at the center of the irradiation field. The shield is a PLA 3D printed case, filled at the bottom with a 3 cm thick layer of aluminum oxide (Al_2_O_3_) powder to stop the electrons and at the top with 1 cm thick layer of 2 mm diameter tungsten (W) sphere aim to stop Bremsstrahlung radiation, minimizing radiation leakage to 0.2%. (E) CBCT image of a mouse positioned in a supine orientation inside the stereotactic mouse positioner, presented in 3 planes: sagittal (top), transverse (middle), and coronal (bottom). Orange arrows indicate the direction of the incident radiation, and the orange highlights indicate the exposed areas in the mouse head; the positioning ensures that eyes are spared from radiation. The thickness of the brain is indicated by a dotted arrow and measured to be 7.6 mm.

**Supplementary Figure 2:**
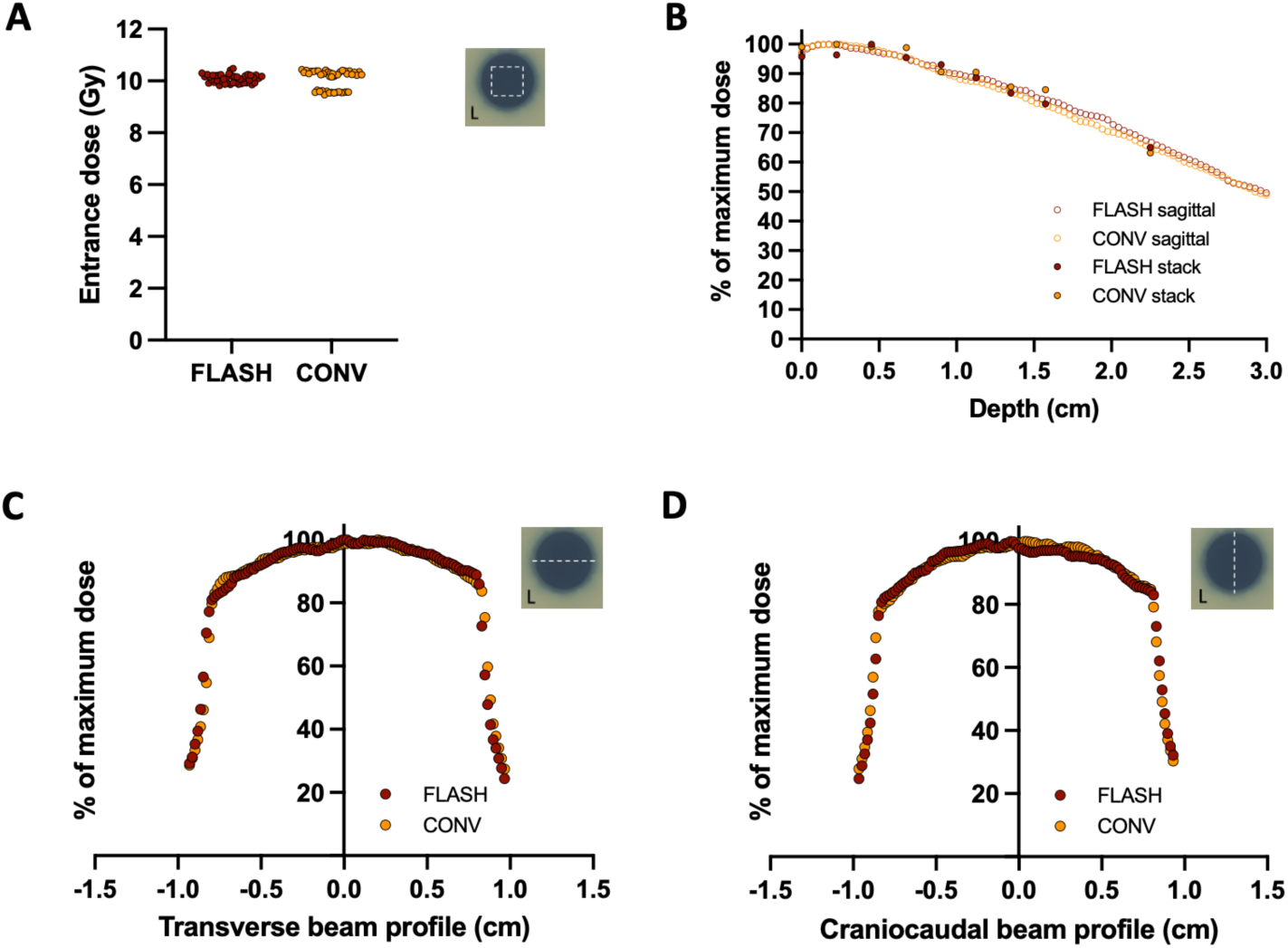
Stanford’s experimental film entrance doses, percentage depth dose and beam profiles for whole brain irradiation. (A) Scan of an experimental radiochromic film (EBT3) used for the measurement of the entrance dose. The square dotted line indicates the area chosen for the analysis. (B) Entrance dose plot showing the doses derived from the experimental radiochromic films from all mice irradiated with FLASH or CONV dose rate. (C) Percentage depth dose curve for the whole brain irradiation beam measured with radiochromic films positioned parallel to the beams path. Black dotted lines indicate the thickness of mouse brain and the absorbed dose at the exit of the radiation beam from the cranial cavity. (D) Transverse and (E) craniocaudal profiles of absorbed entrance dose derived from radiochromic films for both FLASH and CONV dose rate beams. The white dotted line on the exposed films indicates the direction of the profile analysis.

